# Simultaneous analysis of neuroactive compounds in zebrafish

**DOI:** 10.1101/2020.01.01.891432

**Authors:** Douglas Myers-Turnbull, Jack C Taylor, Cole Helsell, Matthew N McCarroll, Chris S Ki, Tia A Tummino, Shreya Ravikumar, Reid Kinser, Leo Gendelev, Rebekah Alexander, Michael J Keiser, David Kokel

**Author notes:** The authors declare no competing interest. Massachusetts Institute of Technology, Cambridge, MA 02139, USA. Genentech, South San Francisco, CA 94080, USA. California University of Science and Medicine, Colton, CA 92324, USA. Paul G. Allen Family Foundation, Seattle, WA 98104, USA.

## Abstract

Neuroactive compounds are crucial tools in drug discovery and neuroscience, but it remains difficult to discover neuroactive compounds with new mechanisms of action. To address this need, researchers have developed mid-throughput phenotype-first approaches using zebrafish. This study introduces an open, non-commercial, and extensible hardware/software platform that captures and analyzes drug-modulated phenotypic responses larval zebrafish. We provide full specifications, computer-aided design (CAD) documents, and source code. Accompanying this study, we are also publicly depositing phenotypic data on 3.9 million animals and 34,000 compounds. The data include a high-replicate benchmark set on 14 compounds, a well-controlled reference set of 648 known neuroactive compounds, 20 specialized reference sets, a library of 1,520 FDA-approved drugs, 3 screening libraries. This open data resource is curated, structured, tied to extensive metadata, and available under a Creative Commons CC-BY license.

## Introduction

Disorders of the central nervous system (CNS) affect 100 million Americans at an economic burden of $920 billion per year (1). Despite this, CNS drug discovery rates have declined (2). Most projects screen for high-affinity interaction with one target (3). Although extremely high-throughput, these screens require prior knowledge of the disease-linked targets, which is limited for CNS disorders (4, 5). Although most projects are target-first, most first-in-class drugs approved by the U.S. Food and Drug Administration (FDA) from 1999–2008 were discovered phenotype-first (6), suggesting that many CNS drug discovery projects would benefit from phenotype-first screens.

In contrast to target-based screens, phenotypic screens rely less on established pathogenesis and can identify compounds with previously unknown and multitarget pharmacological actions. In many historical cases, a drug was discovered first, and its mechanism only later (7, 8). For example, the antidepressant activities of tricyclics and monoamine oxidase inhibitors were discovered in psychiatric hospitals by observing patients. These discoveries implicated serotonin in depression and lent to the development of selective serotonin reuptake inhibitors (SSRIs) (9). Such phenomenological discoveries are responsible for many prototypical neuroactive drugs. Consequently, we and others have sought to scale phenomenological discovery to higher-throughput using animal models for CNS drug discovery.

Zebrafish larvae and embryos have long been used to assay environmental toxicants (10, 11) and as models for vision (12– 16), threat response (17), memory (18), algesia (19–21), and sleep (22–24). These successes in the laboratory have extended to the clinic: In a rare example of bench-to-bedside, the FDA approved lorcaserin as an antiepileptic, based significantly on evidence in zebrafish (25). More recently, a zebrafish model was used in the life-saving treatment of a 12-year-old patient (26). Genetic and compound-induced disease models in zebrafish larvae have shown promising consistency with rodent models (27, 28).

Zebrafish are well-suited for *phenotypic profiling*, a quantitative, high-throughput approach to phenotype-first compound discovery (23, 29). Profiles are quantitative readouts of aggregate animal movements in multiwell plates. These experiments often employ acoustic, photic (light-based), and other stimuli to perturb the animals’ behavior in an effort to reveal more compound-induced behavioral changes. Previous screens identified new neuroactive compounds and predicted their targets, later supported by in vitro assays (23, 30–33). Diverse compounds have been identified, including photoactivatable transient receptor potential channel A1 (TRPA1) ligands (34), antiepileptics (35), antipsychotics (36), appetite modulators (37), and anesthetic-like compounds (38, 39).

One way to predict the pharmacology of a mechanistically novel compound is by association to a compound of known pharmacology. This *guilt-by-association* approach links novel compounds to known ligands, but it requires both *reference profiles* for compounds with known pharmacology and a way to measure similarity between profiles.

Here, we describe the SauronX platform in detail. We benchmark the system in a machine learning approach, first on 14 quality–control (QC) compounds, then on a chemical library of 648 known CNS ligands, all of which we release within a 3.2 million zebrafish, 34,000 compound phenotypic screen public dataset.

## Results

### An open platform enables high-throughput capture of phenotypic data

We sought to develop an open platform for behavioral profiling. We defined 3 criteria: the ability to screen without interruption, reproducibility of analyses, and extensibility to add or remove hardware. We modified an existing system to achieve this (36). The new hardware–software platform has been used to assay N,N-Dimethylaminoisotryptamine (isoDMT) analogs (40), a non-hallucinogenic ibogaine analog with therapeutic potential (41), and toxicants at the U.S. Army Medical Research Institute of Chemical Defense (USAMRICD).

The setup is shown in Fig. 1a. Plates are positioned on a flat translucent stage, fixed in a groove so that sound propagated through the stage contacts the plate uniformly. The plates are illuminated from the bottom with infrared light through an acrylic diffuser and recorded with an overhead camera while light and sound stimuli are applied (Fig. 1a and SI, Fig. S1). The digital camera is mounted to a telecentric lens with an infrared pass filter so that the photic stimuli do not affect the video. The lens eliminates parallax, resulting in all wells having the same apparent dimensions, simplifying feature calculations and eliminating parallax corrections as potential confounding variables. The camera captures 1 Mpx to 6 Mpx (16-bit depth) images at a preset frame rate of 100 Hz to 150 Hz (SI, Movie S1). Nanosecond-resolved timestamps corresponding to the image sensor acquisition are used to precisely synchronize captured frames with stimuli. Computer-aided design (CAD) files (SI, Fig. S1) and related information are available in the Data Repository (42).

**Fig. 1.**
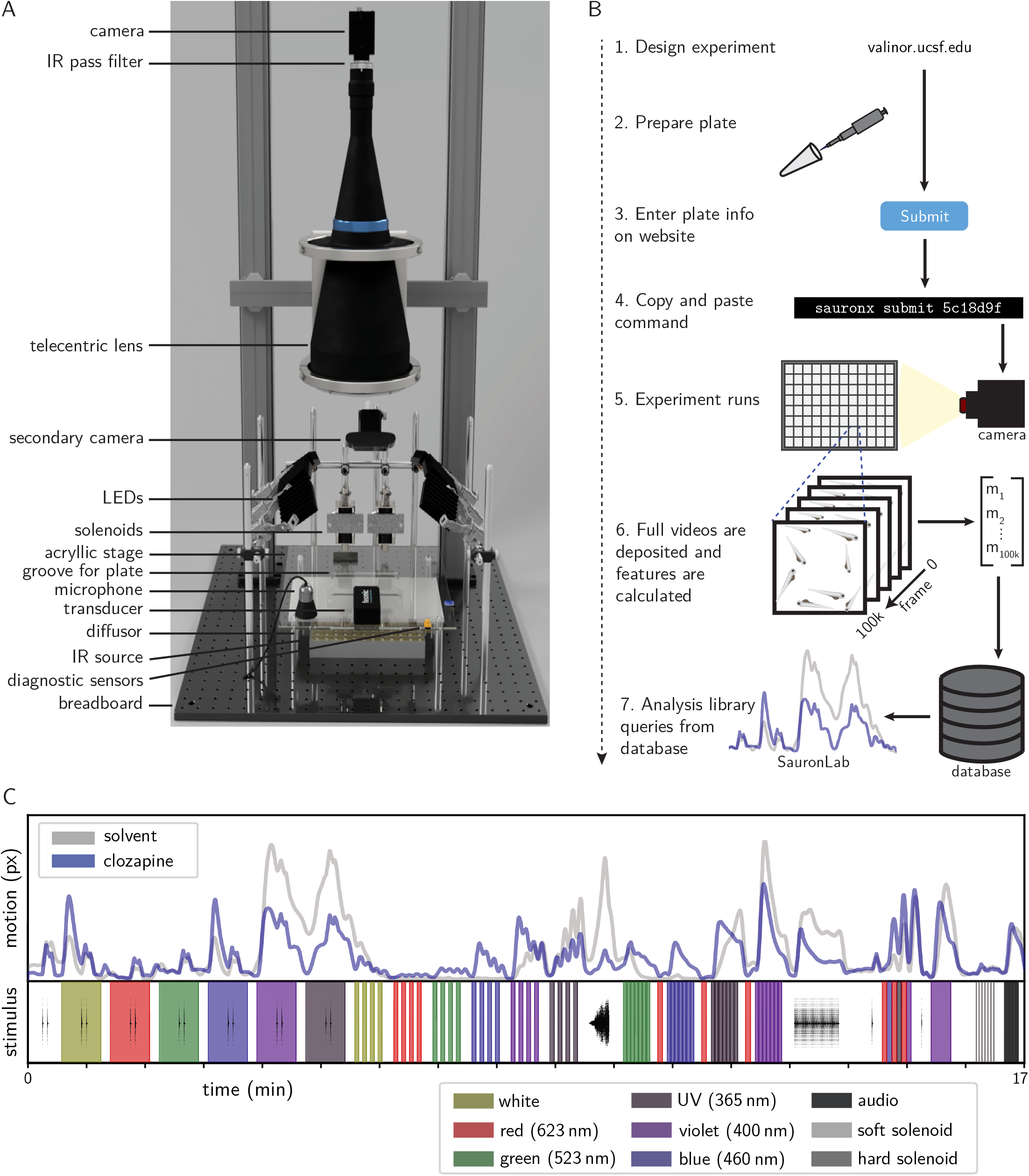
Overview of methods. (**A**) Front view of the instrument. (**B**) Stages of the experimental/computational pipeline. (**C**) Example motion-trace for wells treated with vehicle (dimethyl sulfoxide (DMSO)) or clozapine at 50 µM. Top: motion within the well as a function of time. Bottom: stimuli applied over time. The shaded colors represent high-intensity LED light application, the black lines depict the waveforms of audio assays, and the gray vertical lines (at the end) denote the application of acoustic stimuli by solenoids. N = 12 wells / condition.

To expand the repertoire of observable behavioral responses, stimuli are applied during capture. Photic stimuli are delivered via 6 overhead LED arrays (SI, Table S1). Acoustic stimuli from audio files are delivered through surface transducers mounted on the stage. A microphone, photosensor, and secondary camera verify the delivery and timing of stimuli. These stimuli evoke compound-dependent behaviors that would not otherwise be observed. For example, we observed a compound-modulated ‘step’ response to 355 nm ultraviolet light, which is visible to zebrafish (43). This response differs markedly from 400 nm light (SI, Fig. S3 and SI, Movie S2).

We use a 4-step workflow (Fig. 1b). Animals are anesthetized in cold water and dispensed into the wells of a multiwell plate, dosed, and incubated for 1 hr. The plates are placed in the instrument, and the animals are acclimated in darkness for 5 minutes. A battery of stimuli is then applied while video is recorded. Videos can be analyzed in many ways, including tracking of animals. For the experiments in this manuscript, we used multiple (8) animals per well and calculated a simple feature (*motion-trace*) of aggregate locomotor activity over time (SI, Equation 1). Although using multiple animals per well complicates per-animal tracking, it resulted in much higher algorithm performance (discussed later). Fig. 1c and SI, Fig. S3 show example traces under a standard battery for vehicle (DMSO solvent) or the antipsychotic clozapine.

We use a web interface to design plate layouts, stimulus batteries, and experiments, as well as to organize and search for genetic constructs and compound stocks. A custom language called Gale can be used to design assays from simple expressions (but is not required). The hardware is driven by custom software, which we have released as open source. Post-processing of data is not coupled to capture, allowing many plates to be run in sequence without interruption. After a run completes, the videos are compressed and archived permanently, and data is inserted into a relational database on a remote server.

The database incorporates coarse-grained and fine-grained data. The coarse-grained data, such as hierarchical grouping of experiments, simplifies search. The fine-grained data is included for reproducibility and post-hoc diagnostics. For example, compound treatments are indicated by ‘batch’, with supplier information and lot numbers. In developing the system, we identified information required to conduct reproducible, audit-able analyses. In accordance, we propose a minimum information standard (44) at https://osf.io/nyhpc/.

These data are used in an open source analysis platform (sauronlab), which provides tools for search and analysis. Analyses include quantifying the strength of phenotypes, classifying and clustering phenotypes, analyzing mechanism of actions (MOAs), and searching for similar phenotypes. All analyses are tied to a timestamp that restricts the data queried from the database, ensuring that results do not change when new data is added. Notebooks illustrating these analyses with code and output are available at https://github.com/dmyersturnbull/sauronlab-publication.

In contrast to commercial phenotyping systems, the hardware, data storage, and analysis are uncoupled. Videos are efficiently compressed and can be stored indefinitely and analyzed with additional methods at any point. Although the hardware is larger than most commercial systems at 61 cm×61 cm×114 cm, this simplified construction and enabled rapid iteration between analyzing data and adapting hardware. As part of an effort to develop an open alternative to commercial systems, we benchmarked the platform’s ability to distinguish compound-induced phenotypes.

### Compound-induced phenotypes are reliably detected and distinguished in a benchmark

We wanted to evaluate the platform in a way that is not constrained to a single phenotype. Specifically, we sought to test the ability to detect compound-induced phenotypes (*detection criterion*), identify phenotypes caused by the same compounds while distinguishing those caused by different compounds (*distinction criterion*), and group compounds with similar mechanisms or effects (*grouping criterion*).

First, we curated a set of 14 compounds with diverse structures and MOAs (Table 1 and SI, Table S2). The lethal control used a high dose of the anesthetic eugenol, which is routinely used as a humane method to euthanize fish (45, 46). These 14 compounds and 2 controls formed the QC set. Experiments were run using 7-day-old wild-type zebrafish 1 h post-treatment under a standard battery (Fig. 1c).

**Table 1.**
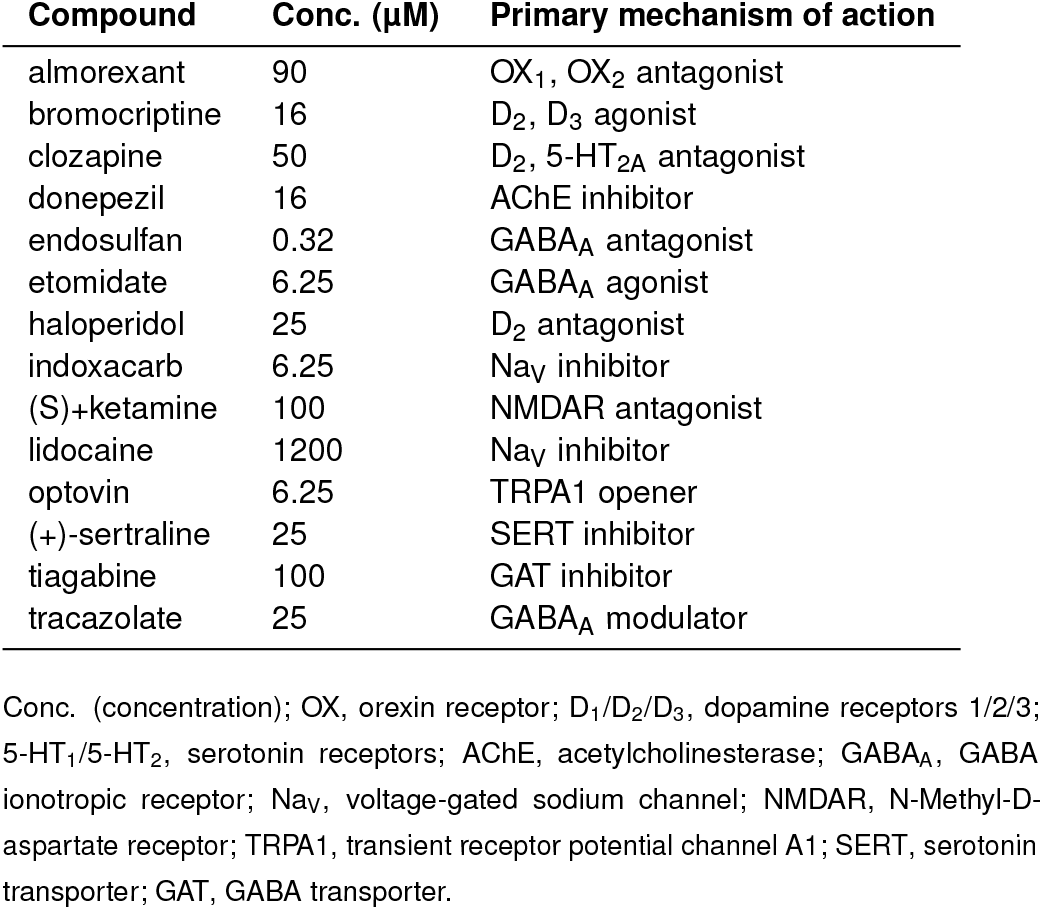
Quality–control compounds with optimal concentrations.

For each compound, we selected a 5-point logarithmic concentration gradient to capture the range between phenotypic inactivity and lethality. 8 replicate plates were screened, with all compounds and concentrations on each plate in random positions. For each compound and concentration, a binary treatment–vehicle Random Forests (RF) classifier was trained to assign the motion vectors as either *treatment* or *vehicle*. The same procedure was used for treatment–lethal models. We plotted the resulting out-of-bag accuracy values in concentration– response curves. Due to the high dimensionality, such curves are not expected to be sigmoidal or even monotonically increasing. For most compounds, treatment–vehicle accuracy increased with concentration, while treatment–lethal accuracy dropped sharply at high concentrations (Fig. 2a, SI, Fig. S2, and SI, Table S3). Notably, treatment–lethal accuracy was high even for sedating doses of the anesthetic etomidate (38), indicating that sedation and lethality were distinguished.

**Fig. 2.**
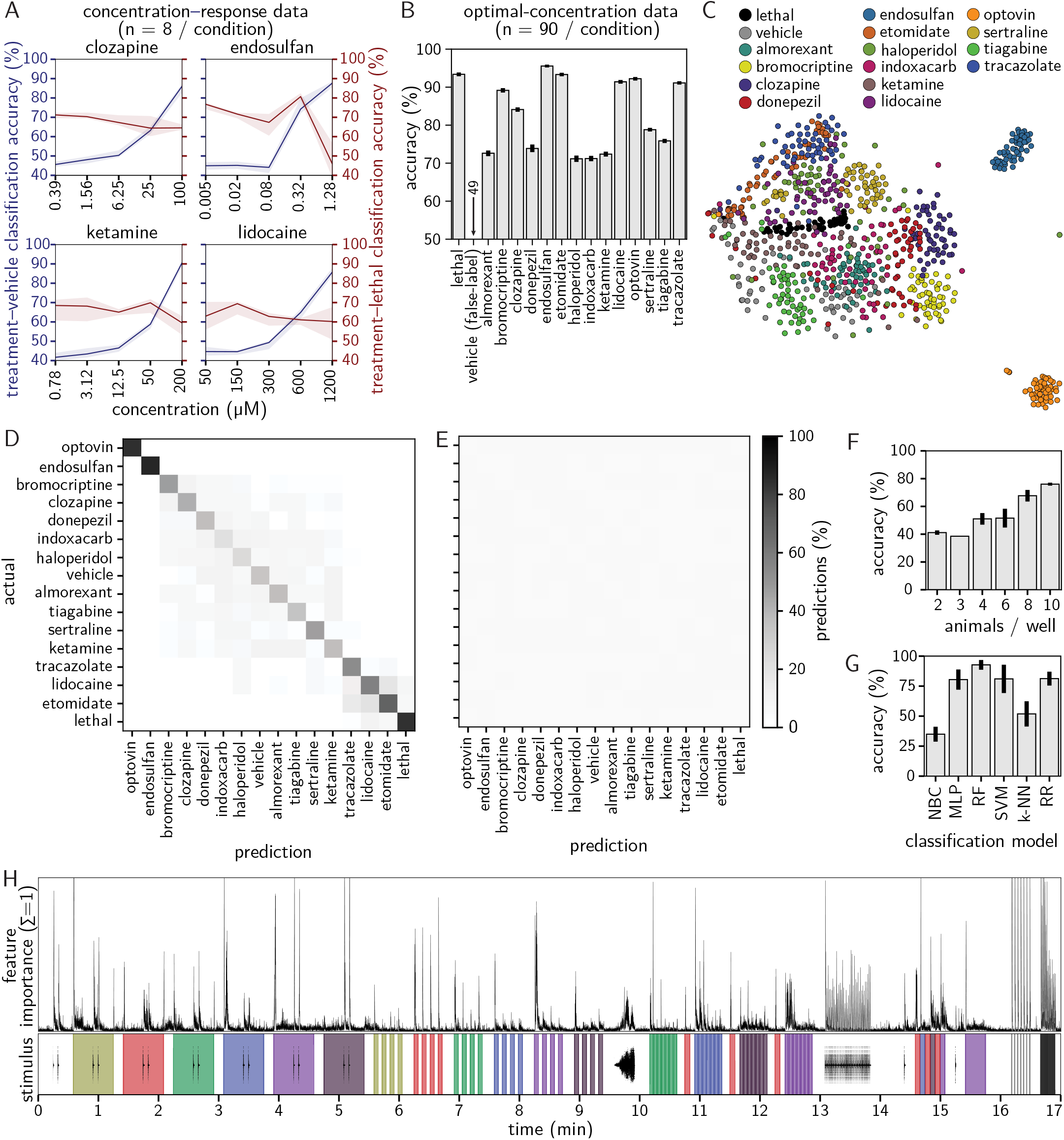
Results for the QC experiments. Abbreviations: NBC, naive Bayes classifier; MLP, multi-layer perceptron; RF, Random Forests; SVM, support vector machine; k-NN, k-nearest neighbor; RR, ridge regression. (**A**) Concentration–response curves for treatment–vehicle (left axis; blue) and treatment–lethal (right axis; red) accuracy. Opaque lines denote the median accuracy, and shaded regions denote a 95th percentile confidence interval by bootstrap. N = 8 wells/condition. (**B**) Treatment–solvent classification accuracy by compound on the optimal-concentration QC set. N = 90 wells/condition. (**C**) T-SNE projection of motion vectors in the optimal-concentration QC set. Each point denotes one well. N = 90 wells/condition. (**D**) Confusion matrix from a multiclass classification model (Random Forests) on the optimal-concentration QC set. N = 90 wells/condition. (**E**) Confusion matrix from a corresponding model trained on false-labeled vehicle-treated wells. N = 18 wells/condition. (**F**) Treatment–treatment accuracy (Random Forests) by number of animals per well in a dedicated experiment. Error bars denote an 80% confidence interval by bootstrap over wells. N = 12 wells/condition. (**H**) Treatment–treatment accuracy as evaluated by different classification algorithms in the optimal-concentration QC set. The extents of the error bars mark the values for the two individual plates. N = 90 wells/condition. NBC, naive Bayes classifier; MLP, multi-layer perceptron; RF, Random Forests; SVM, support vector machine; k-NN, k-nearest neighbor; RR, ridge regression. (**H**) Top: Frame-by-frame feature weights for the optimal-concentration multiclass model (values sum to 1). Extreme values were cropped for visual clarity. Bottom: Stimulus battery as in Fig. 1c. N = 90 wells/condition.

Seeking a way to identify lethal compound treatments that could be interpreted more directly, we trained You Only Look Once v5 (YOLOv5) (47, 48) deep-learning object-detection models. Single frames from 46 wells were annotated by drawing rectangles around individual animals and labeling them *alive* or *deceased*, based on morphology. For potential future applications, we also included other phenotypes, labeling lateral orientation (as a sign of loss of righting reflex) and curvature (as a sign of active motion). Considering only *live*/*deceased, live* were detected with 93% precision (SI, Fig. S4), and *deceased* with 57%. The low precision for *deceased* likely resulted from a relative paucity of training examples (lethal concentrations are preferentially avoided), but an estimate *E*_*d*_ for the number of deceased animals can be counted as *E*_*d*_ = 8 *™ N*_live_.

Using these data, we set an ‘optimal’ concentration per compound by balancing phenotypic strength with non-lethality (Table 1). This *optimal-concentration* set was screened in 15 replicate plates, with 6 replicates of each compound per plate (SI, Fig. S3). In compound–vehicle models, the mean accuracy was 93%. By contrast, a straw man analysis on randomly false-labeled controls yielded 49% for vehicle–vehicle comparisons. This established that compounds could be separated from controls, meeting the detection criterion.

We then visualized the phenotypes together using tdistributed stochastic neighbor embedding (t-SNE) (49). Each compound generated a cloud of replicate profiles generally separate from the controls and other compounds (Fig. 2c), indicating an ability to identify phenotypes caused by the same compounds and distinguish them from others. A RF multiclass classifier was trained to quantify this.

The out-of-bag predictions were visualized in a confusion matrix (Fig. 2d). The labels were sorted by an algorithm that maximized block–diagonal structures, grouping like phenotypes. The diagonal was high (mean=94%), reflecting accurate self-classification and phenotypic uniqueness. The classifier distinguished several compounds, such as almorexant and tiagabine, that were poorly separated in Fig. 2c. As an adversarial experiment to test for shortcut learning or memorization, we collected a dataset of only vehicle-treated wells and false-labeled them to mimic the real dataset. Classifiers were unable to distinguish the false-labeled treatments (Fig. 2e), supporting the distinction criterion.

Grouping of compounds (grouping criterion) was harder to assess with few compounds. Although generally distinguishable, lidocaine, etomidate, and tracazolate were sorted nearby. These compounds reduced movement, analogous to their effects in humans, but they evoked noticeably distinct responses to stimuli (SI, Fig. S3). This offered anecdotal but encouraging support for grouping.

### Benchmarks guided the optimization of experimental protocols and computational methods

We hypothesized that this approach of classification on a QC set served as a general evaluation method to guide experimental design. We applied it to design a stimulus battery, optimize experimental and computational methods, and quantify the impact of potentially confounding variables.

In a data-driven approach to design a battery, we compared 53 different 30 s to 60 s behavioral assays (Tables S4, S5). Assays that provided high classification accuracy were included in the final battery. Background (stimulus-free) assays had notably low performance and most of the acoustic assays with pure tones yielded little information. While pure tones are commonly used (50), acoustic assays generated from complex environmental sounds resulted in higher accuracy. Assays with high-frequency light stimuli and those with simultaneous photic and acoustic stimuli also performed well. Likewise, for the optimal-concentration classifier (Fig. 2d), heavily weighted frames occurred near stimuli (Fig. 2h), directly highlighting their importance. Although these experiments were based on a small set of well-characterized compounds, this data-driven approach eliminated assays that provided low phenotypic information and suggested that complex assays may be more useful for resolving compound-induced phenotypes, in contrast to the stimuli more typically used in such assays.

Next, we evaluated how performance changed under different experimental conditions. Using a separate data set, we evaluated using 3, 4, 6, 8, and 10 animals per well. Accuracy increased with the number of animals (Fig. 2f). These two results showed a trade-off between higher performance and the logistics and ethics of using more animals.

Similar to this optimization of experimental protocols, computational methods could be benchmarked. We benchmarked several classification models, testing across hyperparameter sets (Fig. 2g). Neural networks, random forests, and support vector machines (SVMs) outperformed simple models like linear classifiers and k-nearest neighbors (k-NNs). These experiments pointed to a general procedure to compare and optimize protocols.

Finally, we considered the impact of potentially confounding variables such as time of day and exact treatment duration (deviation from 1 h). The 15 variables we tested were not significantly predictive in this highly controlled regime (SI, Fig. S5). However, well locations can be predictive in experiments that lack positional randomization. In addition, some variables may become predictive if their variance increases; for example, time of day may be predictive in an experiment if data collection continues overnight. We posited that this approach could be used to assess confounding for diverse experimental setups.

### 104 phenotypically active CNS ligands provide reference phenotypic profiles

To predict mechanisms for novel compounds, we collected a set of reference profiles from compounds with diverse pharmacological actions. We used the screen-well Neurotransmitter library (*NT-650*, Enzo Life Sciences), which contained 648 CNS ligands. A fully randomized screen was performed, generating approximately 7 replicates per compound at 33 µM in a 648-compound reference set.

To identify hit compounds, per-compound treatment– vehicle and treatment–lethal classifiers were trained as per the QC set. Visualizing the accuracy values, vehicle–vehicle values were centered near 50% (Fig. 3) as expected. Accordingly, the treatment–vehicle values were long-tailed, indicating that compounds falling within this tail were likely active. To select phenotypically active compounds (hits), we applied an accuracy threshold that excluded 99.5% of vehicle–vehicle comparisons, yielding 106 nonlethal hit compounds. Only 1 compound, tetrahydrodeoxycorticosterone, was lethal at the concentration tested, though other compounds may have been toxic but nonlethal.

**Fig. 3.**
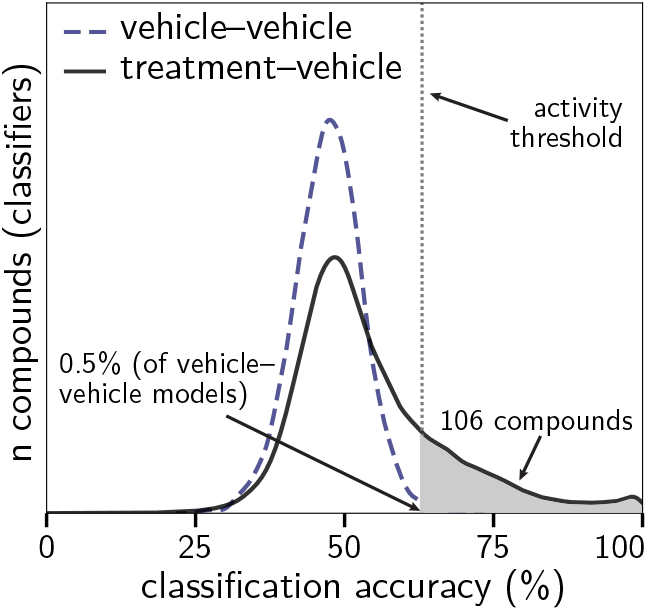
Distribution of prediction accuracy for treatment–vehicle (black, solid) and vehicle–vehicle (blue, dashed) classifiers. The hit threshold is shown at x=63% accuracy. N = 648 non-control compounds (treatment–vehicle). (N = 200 random subsamples for vehicle–vehicle.)

To assess which classes of compounds were more phenotypically active, we grouped compounds by the neurotransmitter systems that they primarily target (Fig. 4). Dopaminergic and serotonergic systems were enriched for phenotypic activity, while adenosinergic, purinergic, and glutamatergic were depleted, although all 13 had at least one hit. Aside from the GABA ionotropic receptor (GABA_A_), enriched protein targets were mostly monoaminergic, including monoamine transporters and dopamine, serotonin, histamine, and muscarinic receptors.

**Fig. 4.**
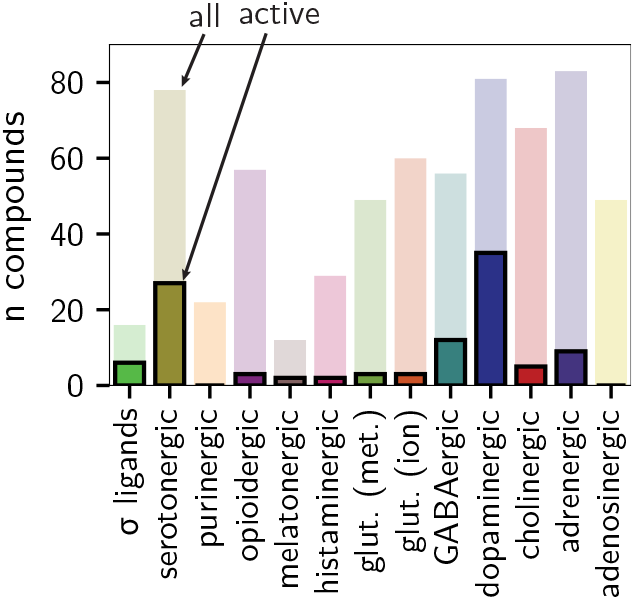
Distribution of hits (opaque) and total compounds (translucent) per major neurotransmitter system. N = 104 compounds. glut. (met.), metabotropic glutamatergic; glut. (ion.), ionotropic glutamatergic.

Multiclass models were then trained to distinguish between the 104 treatments, along with controls. The results were visualized in a sorted confusion matrix (Fig. 5). A strong diagonal indicated that the compounds were phenotypically coherent. Several clusters were observed, including GABA_A_, dopamine transporter (DAT), and dopamine, glutamate, and melatonin receptor ligands. These data indicate that the behavioral screening paradigm is capable of distinguishing neuroactive molecules that interact with discrete neurotransmitter systems. Importantly, multiple chemical scaffolds were present per cluster, indicating scaffold hopping (51) and illustrating the potential for this approach to be used to discover structural starting points for new drugs.

**Fig. 5.**
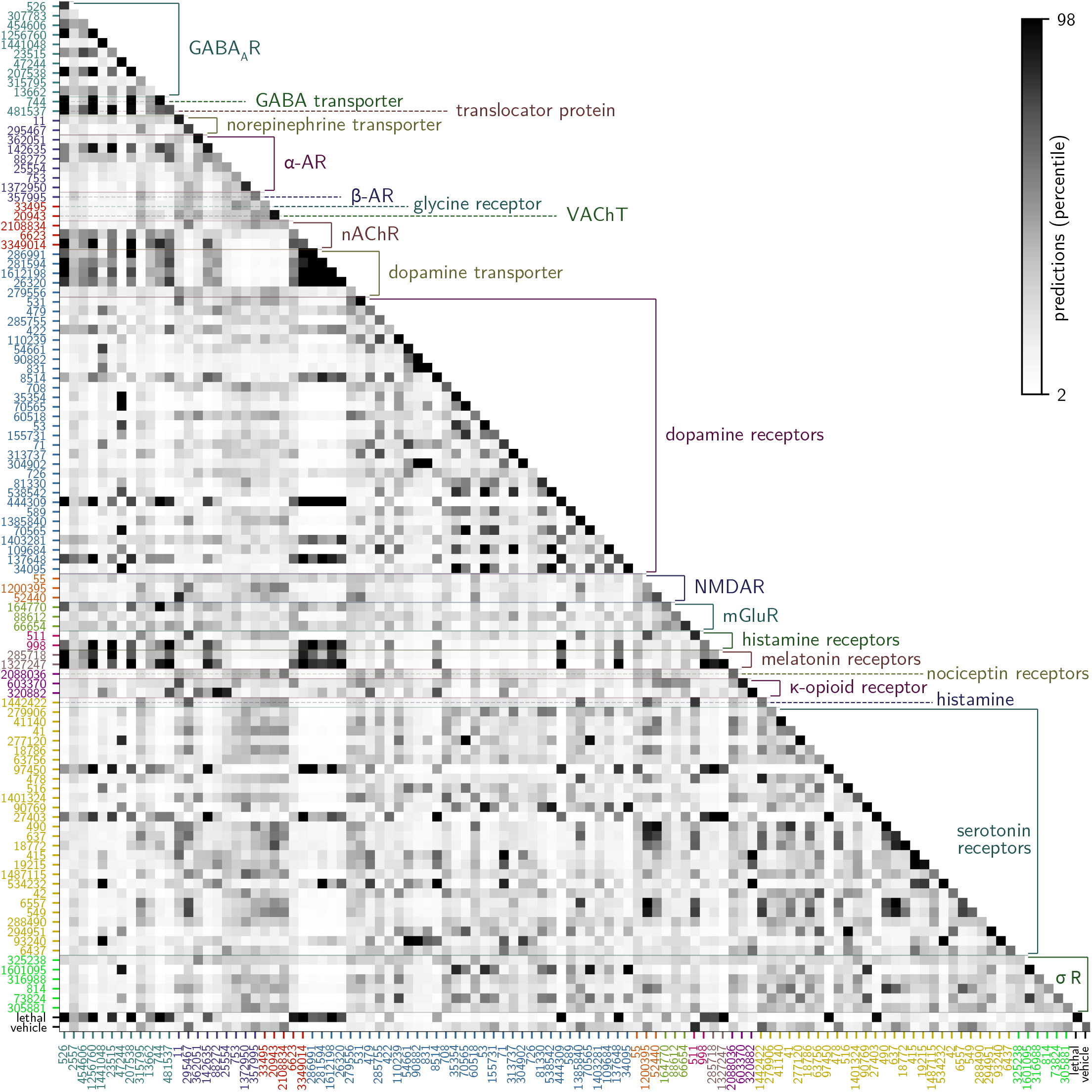
Confusion matrix of the 106 NT-650 hit compounds, plus vehicle and lethal controls. Sorting by MOA targets as provided by Enzo. Range from the 2nd percentile to 98th percentile. Labels are ChEMBL IDs; axis colors indicate the class as per Fig. 4. Arbitrary colors on the diagonal are used to differentiate adjacent labels. N = 530 (compound-treated); 110 (vehicle-treated); 30 (eugenol-treated). GABAA, GABA ionotropic receptor; α-AR, α adrenergic receptor; β-AR, βadrenergic receptor; VAChT, Vesicular acetylcholine transporter; nAChR, nicotinic acetylcholine receptor; NMDAR, N-Methyl-D-aspartate receptor; mGluR, metabotropic glutamate receptor; σ R, σ receptor.

## Discussion

Here we presented an open platform for behavioral phenotyping in zebrafish and posted complete specifications. To facilitate data mining, we publicly deposited phenotypic data for 34,000 compounds and 3.2 million animals. Focusing on a high-replicate, well-characterized quality-control subset of this data repository, we found that machine learning classifiers can readily detect, distinguish, and group known drugs using their behavioral profiles.

Prior studies validate behavioral profiling as a way to discover and characterize neuroactive compounds. Different hardware, zebrafish strains, and computational methods have been applied, and this diversity calls for quantitative evaluations. We found that classification in a quality–control set provided an intuitive and powerful metric to summarize performance. This approach has immediate applications, such as optimizing protocols and assessing the impact of confounding variables. In particular, positional confounding can significantly affect results, supporting a need for treatment randomization. The screen-well Neurotransmitter library provides compounds physically arranged by their major pathways, illustrating how this confounding could solely explain a promising result. We provide a lower bound on performance and hope this will invite comparisons using the same approach or the development of superior or complimentary benchmarks.

Certain modifications could expand the observable subset of compound-induced movement behaviors. First, we used a concentration of 33 µM for the NT-650 screen, but the concentration–response experiments indicated that some compounds were phenotypically inactive below 100 µM. Second, affecting complex states such as aggression, addiction, or learning may improve resolution. We used a trivial readout for high-dimensional movement behaviors, but tracking (52–54), optical flow (55), probabilistic models (56), and deep learning (57) have been successful in analyzing similar data.

Finally, technologies like RNA-seq and mass spectrometry could be applied in concert with behavioral experiments as powerful, high-throughput, and high-dimensional approaches to delineate the mechanisms underlying behavioral modifications. Future studies will likely leverage advances in many of these areas to improve the resolution of behavioral profiling.

With the release of full open hardware and software platform specifications, along with an accompanying data repository of 34,000 compounds on 3.2 million zebrafish spanning 7 years, we hope to spur analysis and development opportunities of open systems for behavioral phenotyping and the discovery of novel neuroactive compounds.

## Methods

### Animal husbandry

Zebrafish husbandry was as described (58). Embryos were from group matings of wild-type zebrafish (Singapore, ZFIN:ZDB-GENO-980210-24) raised on a 14/10-hour light/dark cycle in 28 °C egg water (Instant Ocean (003746) with NaHCO_3_ to pH 7.0–7.4) (58) until 7 dpf. Animals were maintained in a facility accredited by the Association for Assessment and Accreditation of Laboratory Animal Care (AAALAC). Experiments were performed in accordance with protocols approved by UCSF’s Institutional Animal Care Use Committee (IACUC) and in accordance with the Guide for the Care and Use of Laboratory Animals (59).

### Software and data availability

Hardware information, protocols, links to software repositories, extended supplemental data, and the full database are available at https://osf.io/3dp6x. Software is released under an Apache 2.0 license.

### Instrument

Components used in these analyses are documented in SI, Section A. Some were replaced afterward; current recommendations are at https://osf.io/3dp6x. A PointGrey Grasshopper GS3-U3-41C6M-C camera (FLIR Integrated Imaging Solutions) and infrared pass filter were used (LE8744 polyester #87, LEE Filters). Six high-intensity LED arrays were positioned overhead, with 4 LEDs per array (Osram Sylvania; LED Engin; New Energy). Two surface transducers were fastened on the stage (5 W transducer, Generic) and used with a 150 W amplifier (APA150, Dayton Audio). Two 36 V push–pull solenoids (SparkFun Electronics) were positioned near the top of the plate, one contacting the stage directly, and the other contacting a 1 mm-deep strip of synthetic felt. Audio files (SI, Sound Files S1–3) are provided along with sound pressure level (SPL) measurements (SI, Table S6).

An Arduino Mega 2560 rev 3 (Arduino.cc) drove the LEDs, solenoids, and small sensors while a computer directly controlled the microphone, transducers, and camera. The camera streamed raw data to a high-performance drive without buffering. Videos were 1600 × 1068 in 8-bit grayscale. Videos were trimmed, compressed with High-Efficiency Video Encoding (HEVC) using Constant Quantization Parameter (CQP) 15 and partitioned into region of interests (ROIs) for wells (SI, Section B).

### Data collection and filtration

Healthy larvae were sorted and then immobilized with cold egg water with 25 mL of 4 °C added to 12 mL room-temperature egg water in a 100 cm petri dish containing about 1,000 fish. 8 larvae in 300 µL were then distributed by pipette into the wells of 96-well plates, using trimmed tips to avoid injuring the animals. Plates were incubated at room temperature for 1 hr, at which animals were mobile.

For QC experiments (SI, Table S2), compound plates and aliquots were stored at ™20 °C. 2.0 µL of solvent-dissolved compound was then added to each well. Solvents were dimethyl sulfoxide (DMSO) except for donepezil (water). Some donepezil wells had less than 2.0 µL remaining due to evaporation (annotated in the database). Each concentration–response curve included 5 concentrations on a logarithmic scale with an additional hypothesized ideal concentration (SI, Table S3). The optimal-concentration QC set was replicated across 15 plates, applying 6 replicates of the 14 compounds and 2 controls (16 × 6 = 96). The vehicle-only adversarial control experiment was collected with 3 plates using earlier hardware and a different battery. However, optimal-concentration QC accuracy was high when subsampled to 3 plates. 5/14 optimal-concentration plates and 1/9 concentration–response plates were excluded because hardware sensors flagged them for potential problems.

The screen-well Neurotransmitter library (Enzo Life Sciences) was purchased in solution at 10 mM (peptides 100 µM) in 2015 and stored at ™80 °C. A Biomek FX^P^ (Beckman Coulter) was used for randomization.

For NT-650, 1 µL was added per well to yield 33 µM, except for peptides at 0.33 µM. Treatments were randomized across plates and wells. Each plate contained 14 DMSO, 8 water, and 6 lethal eugenol controls, except for 1 of every 7 plates due to an uneven split. 7 replicates were screened per compound, with deviation from 7 due to a subsequent filtration. 13/80 plates were excluded based on sensor readout. We also filtered 23/7680 wells that had insufficient volume of compound in the daughter plate (SI, Table S7).

### Phenotype analysis

Pre-interpolation motion vectors were:

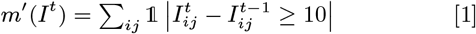

Where *I*^*t*^ is the image matrix at 1-indexed frame *t*. The threshold 10 was chosen by comparing a histogram of pixel intensity changes in wells with and without fish. The final motion *m* was then quantified by linear interpolation of *m*^*′*^ values and image sensor acquisition timestamps to align frames and stimuli.

RFs were trained with scikit-learn 0.24.1 (RRID:SCR_002577) (60) with default hyperparameters except for the number of estimators, which was 20,000 for treatment–vehicle and 40,000 for multiclass classification. Reported accuracies were out-of-bag. YOLOv5 (Git tag v5.0; commit `f5b8f7d54c9f`) models were trained on images at time 16:55 from 11 plates, 46 wells, and 338 animals. Boxes were drawn around animals and labeled using labelImg. Set sizes were: 191 *alive*, 61 *deceased* (training); and 59 *59*, 27 *deceased* (training). Augmented images were generated under *D*_4_ symmetry operations. Cross-validation was performed with a 3:1 train:test split.

For NT-650 hit-calling, 4 replicate treatment–vehicle classifiers were trained per compound. Per classifier, all replicate treatment wells were compared with the same number of randomly sampled vehicle wells, restricted to the plates containing the compound treatment and with the same solvent (DMSO or water). Amoxapine (CHEMBL1113) was dissolved in N-methyl-2-pyrrolidone (NMP); it was compared to DMSO. Lethality was detected by an analogous procedure. For the NT-650 multiclass problem, the mean was taken over 5 confusion matrices, each trained on a stratified subset with 4 wells per compound.

### Visualization

Motion-trace visualizations were smoothed from 100 Hz to 10 Hz with a sliding window. T-SNE parameters were scikit-learn defaults. Concentration– response curves were computed with 1,000 bootstrap samples. kernel density estimate (KDE) were Gaussian, calculated with statsmodels 0.10 (RRID:SCR_016074) (61): `kdensityfft(kernel=gau, bw=normal_reference`. Matrix sorting used confusion matrix ordering (CMO) (62) via clana version 4.0; `simulated_annealing` was called with default arguments.

### Control experiments and battery design

Assays subject to constraints were generated exhaustively, using LED assays, pure tones and environmental sounds, and combinations. Assays were ranked by the 80th percentile of their accuracy over the 16 unique treatments. The number-of-fish experiment used 2 randomized plates, 2 plates / condition. The strain comparison used 2 plates / strain.

## ACKNOWLEDGMENTS

The authors thank Louie Ramos, Julian Castaneda, Veronica Manzo, Nolan Wong, Madison Burns, Vy Nguyen, and Alexis Marquez for animal husbandry; Giancarlo Bruni for hardware; Capria Rinaldi and Ashley Oh for referenced data; Steven Chen, Michelle Arkin, Adam Renslo for Biomek access and handling; the Dow Chemical Company and Robert Tombari for providing compounds; and Eric Lam for fabrication assistance. Funding was provided by the National Institute on Alcohol Abuse and Alcoholism; the Paul G. Allen Family Foundation, the Genentech Fellowship Program; National Institutes of Health grant 4T32GM6754714; and grant number 2018-191905 from the Chan Zuckerberg Initiative Donor-Advised Fund at the Silicon Valley Community Foundation.

