## Supplementary material for "Simultaneous analysis of neuroactive compounds in zebrafish": supplemental information.pdf

#### This PDF file includes:

Supplementary text
Figs. S1 to S7
Tables S1 to S7
Legends for Movies S1 to S2
SI References

#### Other supplementary materials for this manuscript include the following:

Movies S1 to S2
Audio Files S1 to S3

### Supporting Information Text

#### Hardware and drivers

This section describes the pipeline that was presented in the main text, except for the false-labeled quality–control (QC) adversarial experiment. The current version is run in Linux (Ubuntu 19.10), but some data were acquired in Windows 10. Three instruments were constructed, but only one was used for these data. Data included in the data repository were from these instruments along with legacy instruments.

Four sensors were used for diagnostics: An H2a Hydrophone (Aquarian Hydrophones) with a rubber contact adapter on the stage surface; an overhead Logitech 1080p C930e webcam (Logitech); and a photoresistor and thermistor (Tinker Kit, SparkFun Electronics) under the stage. Sensors recorded for the full duration, padded by 1 s. Hardware and software settings were listed in per-instrument configuration files, which were occasionally modified; these changes were tracked in the database, with each run associated with a particular version. Configuration settings included the region of interest (ROI), framerate, exposure, pin mapping, directory structure, and sensor configuration.

#### Video acquisition and processing

Video was streamed to high-performance M.2 SSD via USB 3.1 Gen 1 (Samsung EVO 970 PRO M.2 1 Tb, Samsung). Both requirements were found to be needed to avoid throttling the acquisition.

Video data was streamed from the camera to the Non-Volatile Memory Host Controller Interface Specification (NVMe) drive in raw image sensor format ('RAW') using custom C++ driver code based on the Spinnaker SDK version 1 (FLIR Systems). After acquisition, timestamps from the firmware clock corresponding to image sensor acquisition were mapped to a `std::chrono::high_resolution_clock` system clock in the drivers. These were then used to trim the video frames to the exact start and end of the battery using timestamps from the image sensor.

Raw frames were then converted to High-Efficiency Video Encoding (HEVC) via ffmpeg 4 using Quick Sync Video (Intel) hardware encoding on an Intel i7-9700K Coffee Lake 8-Core 3.6 GHz processor. This was found to provide better performance with no perceptible loss in quality over both software encoding and hardware encoding on a GeForce GTX 1060 (NVIDIA) graphics card. (The graphics card was then removed because it interfered with using Intel's hardware encoder.)

The exact ffmpeg command used was:

```
ffmpeg \
-loglevel warning \
-hwaccel image2 \
-nostdin \
-c:v rawvideo \
-framerate 100 \
-video_size ${width}x${height} \
-pixel_format gray \
-start_number ${first_frame} \
-i "${path}/%08d.raw" \
-y \
-c:v hevc_qsv \
-g 100 \
-q:v 15 \
-preset veryfast \
-load_plugin 2 \
-r 100 \
"${path}/qc15.mkv"
```

These parameters were selected after extensive testing; significant compression artifacts were not observed for these values. A typical video was circa 200 Gb in image sensor output (RAW files) and circa 5 Gb of compressed data for a 17-minute video. Compression ratios were noticeably higher for videos with less motion.

#### Motion interpolation

Preliminary motion-traces were then computed using a custom Scala function (`CdplusFeature(tau=10).apply`), streaming video frames via ffmpeg. To keep the number of features equal to the number of frames, the calculation was:

$$m'(I^t) = \begin{cases} \sum_{ij} \mathbb{1} \left| I^t_{ij} - I^{t-1}_{ij} \geq 10 \right| & t \geq 2 \\ 0 & t = 1 \end{cases} \quad [1]$$

These values were inserted into the database. The motion vectors were then interpolated in Sauronlab using the acquisition and stimulus timestamps. The Scipy 1.3.0 (1) function `interp1d` was called with `kind=previous` and `fill_value=extrapolate`.

**Table S1. LED stimuli.**

| Color | Chromaticity | Intensity | Manufacturer | Part number |
|---|---|---|---|---|
| red | 625 nm | $4.6{\times}10^2$ lm | Osram Sylvania | LZ4-40R108-0000 |
| green | 525 nm | $6.6{\times}10^2$ lm | Osram Sylvania | LZ4-40G108-0000 |
| blue | 650 nm | $3.9{\times}10^3$ mW | Osram Sylvania | LZ4-40B208-0000 |
| violet | 400 nm | $3.0{\times}10^3$ mW | LED Engin | LZ4-40UB00-00U7 |
| UV | 365 nm | $8.8{\times}10^2$ mW | New Energy | LST1-01G01-UV01-00 |
| white | 4,000 K | $1.6{\times}10^3$ lm | New Energy | XHP70A-00-0000-0D0BN240E-SB01 |

'Intensity' values are (total) radiant or luminous flux reported in documentation, not measured.

**Table S2. Full information on quality–control compounds.**

| Compound | Concentration (µM) | Primary mechanism of action | Supplier | Catalog |
|---|---|---|---|---|
| almorexant HCl | 90 | orexin receptor 1, orexin receptor 2 antagonist | Selleckchem | S2160 |
| bromocriptine mesylate | 16 | dopamine 2 receptor, dopamine 3 receptor agnoist | Tocris | 0427 |
| clozapine | 50 | dopamine 2 receptor, serotonin 2A receptor antagonist | Sigma Aldrich | C6305 |
| donepezil HCl | 16 | acetylcholinesterase inhibitor | Tocris | 4385 |
| endosulfan | 0.32 | GABA ionotropic receptor antagonist | Sigma Aldrich | 32015 |
| (R)-etomidate | 6.25 | GABA ionotropic receptor agonist | Tocris | 1471 |
| haloperidol | 25 | dopamine 2 receptor antagnoist | Sigma Aldrich | H1512 |
| indoxacarb | 6.25 | voltage-gated sodium channel inhibitor | Sigma Aldrich | 33969 |
| (S)+ketamine HCl | 100 | N-Methyl-D-aspartate receptor antagonist | Sigma Aldrich | 33795-24-3 |
| lidocaine HCl monohydrate | 1200 | voltage-gated sodium channel inhibitor | Sigma Aldrich | L5647 |
| optovin | 6.25 | transient receptor potential channel A1 opener | Hit2Lead | 6197309 |
| (+)-sertraline HCl | 25 | serotonin transporter inhibitor | Sigma Aldrich | S6319 |
| tiagabine HCl | 100 | GABA transporter inhibitor | Abcam Biochemicals | ab120237 |
| tracazolate HCl | 25 | GABA ionotropic receptor modulator | Tocris | 56967 |

OX, orexin receptor; $D_1/D_2/D_3$, dopamine receptors 1/2/3; 5-$HT_1$/5-$HT_2$, serotonin receptors; AChE, acetylcholinesterase; $GABA_A$, GABA ionotropic receptor; $Na_V$, voltage-gated sodium channel; NMDAR, N-Methyl-D-aspartate receptor; TRPA1, transient receptor potential channel A1; SERT, serotonin transporter; GAT, GABA transporter.


Douglas Myers-Turnbull, Jack C Taylor, Cole Helsell, Matthew N McCarroll, Chris S Ki, Tia A Tummino, Shreya Ravikumar, Reid Kinser, Leo Gendelev, Rebekah Alexander, Michael J Keiser, David Kokel

**Table S3. Treatment–vehicle and treatment–lethal concentration–response values.**

| name | C1 | C2 | C3 | C4 | C5 | C6 | V1 | V2 | V3 | V4 | V5 | V6 | L1 | L2 | L3 | L4 | L5 | L6 |
|---|---|---|---|---|---|---|---|---|---|---|---|---|---|---|---|---|---|---|
| almorexant | 1.4 | 3.62 | 22.5 | 50 | 90 | 360 | 86 | 86 | 89 | 90 | 89 | 95 | 97 | 97 | 97 | 95 | 95 | 90 |
| bromocriptine | 1 | 4 | 16 | 50 | 64 | 256 | 86 | 87 | 97 | 89 | 87 | 98 | 98 | 95 | 98 | 92 | 95 | 94 |
| clozapine | 0.39 | 1.56 | 6.25 | 25 | 50 | 100 | 86 | 86 | 86 | 87 | 92 | 95 | 98 | 98 | 97 | 97 | 97 | 97 |
| donepezil | 1 | 4 | 16 | 50 | 64 | 256 | 86 | 86 | 92 | 86 | 87 | 92 | 97 | 98 | 98 | 98 | 98 | 98 |
| endosulfan | 0.005 | 0.02 | 0.08 | 0.195 | 0.32 | 1.28 | 86 | 86 | 86 | 98 | 95 | 92 | 97 | 97 | 97 | 100 | 98 | 92 |
| etomidate | 0.02 | 0.2 | 2 | 6.25 | 20 | 200 | 86 | 86 | 87 | 97 | 100 | 98 | 97 | 97 | 97 | 90 | 90 | 94 |
| haloperidol | 0.39 | 1.56 | 6.25 | 6.25 | 25 | 100 | 86 | 86 | 90 | 90 | 86 | 90 | 97 | 95 | 99 | 99 | 98 | 94 |
| indoxacarb | 0.08 | 0.8 | 6.25 | 8 | 80 | 800 | 86 | 86 | 90 | 86 | 87 | 95 | 97 | 97 | 97 | 94 | 95 | 94 |
| ketamine | 0.78 | 3.12 | 12.5 | 50 | 100 | 200 | 86 | 86 | 86 | 86 | 86 | 98 | 98 | 95 | 98 | 97 | 97 | 95 |
| lidocaine | 50 | 150 | 200 | 300 | 600 | 1200 | 86 | 86 | 86 | 86 | 86 | 94 | 94 | 97 | 97 | 97 | 97 | 94 |
| optovin | 0.01 | 0.15 | 2.25 | 6.25 | 33.8 | 506 | 86 | 86 | 86 | 97 | 95 | 94 | 97 | 97 | 98 | 98 | 98 | 98 |
| sertraline | 0.39 | 1.56 | 6.25 | 12.5 | 25 | 100 | 86 | 86 | 84 | 94 | 94 | 100 | 95 | 95 | 97 | 98 | 95 | 86 |
| tiagabine | 0.48 | 2.4 | 12 | 60 | 100 | 300 | 86 | 86 | 86 | 86 | 92 | 89 | 97 | 97 | 95 | 95 | 95 | 94 |
| tracazolate | 0.05 | 0.35 | 2.5 | 17 | 25 | 120 | 86 | 86 | 86 | 100 | 100 | 94 | 95 | 95 | 95 | 86 | 87 | 97 |

5 values along a log scale and 1 hypothesized optimal value. In some cases, the hypothesized concentration was equal to a log-scale value. C1–6, Concentrations in micromolar; V1–6, out-of-bag accuracy (%) of treatment–solvent models for the corresponding concentrations; L1–6, accuracy in treatment–lethal models.

**Table S4. Performance of 53 behavioral assays.**

| # | Assay | $Q_{80}$[1] | $Q_{80}$[2] | $Q_{50}$[1] | $Q_{50}$[2] | $Q_{100}$[1] | $Q_{100}$[2] |
|---|---|---|---|---|---|---|---|
| 1 | audio :: breath 2x | 20 | 17 | 18 | 13 | 29 | 20 |
| 2 | audio :: drums (2 copies) | 18 | 14 | 10 | 10 | 20 | 19 |
| 3 | audio :: explosion 2x | 14 | 18 | 12 | 13 | 22 | 32 |
| 4 | audio :: ring 2x | 19 | 18 | 16 | 12 | 47 | 38 |
| 5 | audio :: thump 2x | 20 | 19 | 15 | 14 | 28 | 22 |
| 6 | audio+light :: ring under blue | 40 | 26 | 20 | 18 | 47 | 35 |
| 7 | audio+light :: ring under dark | 19 | 20 | 13 | 15 | 34 | 36 |
| 8 | audio+light :: ring under green | 26 | 31 | 14 | 19 | 37 | 37 |
| 9 | audio+light :: ring under red | 24 | 27 | 17 | 18 | 31 | 36 |
| 10 | audio+light :: ring under uv | 26 | 26 | 20 | 21 | 32 | 35 |
| 11 | audio+light :: ring under violet | 25 | 23 | 17 | 17 | 29 | 37 |
| 12 | audio+light :: ring under white | 27 | 22 | 16 | 18 | 32 | 42 |
| 13 | background :: 10s (52 copies) | 20 | 26 | 9 | 15 | 47 | 45 |
| 14 | background :: 30s (2 copies) | 14 | 11 | 8 | 8 | 34 | 21 |
| 15 | light :: blue 1s:1s | 17 | 18 | 11 | 14 | 32 | 32 |
| 16 | light :: blue 2s:2s | 19 | 17 | 14 | 12 | 40 | 34 |
| 17 | light :: blue 3500ms:500ms | 13 | 19 | 9 | 12 | 37 | 38 |
| 18 | light :: blue 4s:4s | 14 | 17 | 10 | 10 | 42 | 35 |
| 19 | light :: blue 500ms:3500s | 18 | 15 | 12 | 12 | 38 | 33 |
| 20 | light :: dark flash :: 1s (60s assay) | 15 | 15 | 9 | 11 | 34 | 34 |
| 21 | light :: green 1s:1s | 17 | 16 | 11 | 10 | 54 | 51 |
| 22 | light :: green 2s:2s | 13 | 14 | 9 | 9 | 54 | 59 |
| 23 | light :: green 3500ms:500ms | 29 | 15 | 22 | 9 | 44 | 51 |
| 24 | light :: green 4s:4s | 12 | 19 | 9 | 11 | 38 | 57 |
| 25 | light :: green 500ms:3500ms | 19 | 15 | 11 | 11 | 48 | 48 |
| 26 | light :: red 1s:1s | 17 | 15 | 10 | 12 | 30 | 28 |
| 27 | light :: red 2s:2s | 13 | 17 | 10 | 10 | 30 | 49 |
| 28 | light :: red 3500ms:500ms | 16 | 15 | 9 | 12 | 36 | 38 |
| 29 | light :: red 4s:4s | 13 | 17 | 9 | 9 | 38 | 44 |
| 30 | light :: red 500ms:3500ms | 13 | 18 | 9 | 10 | 29 | 38 |
| 31 | light :: solid blue :: 30s | 20 | 15 | 13 | 11 | 40 | 35 |
| 32 | light :: solid green :: 30s | 14 | 16 | 11 | 11 | 43 | 37 |
| 33 | light :: solid red :: 30s | 18 | 16 | 13 | 11 | 53 | 51 |
| 34 | light :: solid uv :: 30s | 20 | 18 | 10 | 13 | 35 | 44 |
| 35 | light :: solid violet :: 30s | 24 | 18 | 15 | 13 | 38 | 45 |
| 36 | light :: solid white :: 30s | 20 | 14 | 11 | 11 | 42 | 34 |
| 37 | light :: uv 1s:1s | 19 | 19 | 14 | 14 | 35 | 32 |
| 38 | light :: uv 2s:2s | 19 | 24 | 14 | 17 | 34 | 35 |
| 39 | light :: uv 3500ms:500ms | 24 | 17 | 21 | 13 | 35 | 38 |
| 40 | light :: uv 4s:4s | 23 | 23 | 15 | 17 | 38 | 43 |
| 41 | light :: uv 500ms:3500ms | 18 | 22 | 13 | 15 | 35 | 31 |
| 42 | light :: violet 1s:1s | 28 | 17 | 17 | 13 | 40 | 42 |
| 43 | light :: violet 2s:2s | 21 | 25 | 16 | 17 | 43 | 41 |
| 44 | light :: violet 3500ms:500ms | 20 | 18 | 16 | 13 | 41 | 33 |
| 45 | light :: violet 4s:4s | 25 | 17 | 18 | 14 | 38 | 40 |
| 46 | light :: violet 500ms:3500ms | 18 | 19 | 12 | 13 | 34 | 24 |
| 47 | light :: white 1s:1s | 22 | 17 | 16 | 14 | 53 | 53 |
| 48 | light :: white 2s:2s | 17 | 15 | 10 | 12 | 46 | 46 |
| 49 | light :: white 3500ms:500ms | 15 | 16 | 11 | 10 | 49 | 29 |
| 50 | light :: white 4s:4s | 16 | 21 | 12 | 13 | 43 | 39 |
| 51 | light :: white 500ms:3500ms | 16 | 18 | 11 | 13 | 50 | 50 |
| 52 | taps :: soft 1s | 19 | 16 | 11 | 10 | 27 | 20 |
| 53 | taps :: soft 2s | 21 | 15 | 11 | 12 | 29 | 30 |

Comparison of 53 behavioral assays tested following initial development and assessment.

**Table S5. Final assays in the standard battery.**

| # | Assay | Description | Start | End | Duration |
|---|---|---|---|---|---|
| 0 | 10s background | no stimuli | 0ms | 0:10 | 0:10 |
| 1 | audio :: ring dark | ring sound (1.2s; 2x) | 0:10 | 0:30 | 0:20 |
| 2 | light+audio :: ring white | ring sound (1.2s; 2x) under solid white light (40 s) | 0:30 | 1:20 | 0:50 |
| 3 | light+audio :: ring red | ring sound (1.2s; 2x) under solid red light (40 s) | 1:20 | 2:10 | 0:50 |
| 4 | light+audio :: ring green | ring sound (1.2s; 2x) under solid green light (40 s) | 2:10 | 3:00 | 0:50 |
| 5 | light+audio :: ring blue | ring sound (1.2s; 2x) under solid blue light (40 s) | 3:00 | 3:50 | 0:50 |
| 6 | light+audio :: ring violet | ring sound (1.2s; 2x) under solid violet light (40 s) | 3:50 | 4:40 | 0:50 |
| 7 | light+audio :: ring UV | ring sound (1.2s; 2x) under solid UV light (40 s) | 4:40 | 5:30 | 0:50 |
| 8 | light :: white pulses | 4s:4s pulses of white light (4x) | 5:30 | 6:10 | 0:40 |
| 9 | light :: red pulses | 4s:4s on:off pulses of red light (4x) | 6:10 | 6:50 | 0:40 |
| 10 | light :: green pulses | 4s:4s on:off pulses of green light (4x) | 6:50 | 7:30 | 0:40 |
| 11 | light :: blue pulses | 4s:4s on:off pulses of blue light (4x) | 7:30 | 8:10 | 0:40 |
| 12 | light :: violet pulses | 4s:4s on:off pulses of violet light (4x) | 8:10 | 8:50 | 0:40 |
| 13 | light :: UV pulses | 4s:4s on:off pulses of UV light (4x) | 8:50 | 9:30 | 0:40 |
| 14 | audio :: drums | drums with increasing intensity and pitch (20.073 s) | 9:30 | 10:05 | 0:35 |
| 15 | light :: green flicker | 3.5s : 0.5s on:off green light | 10:05 | 10:45 | 0:40 |
| 16 | light :: red flash | solid red light (5 s) | 10:45 | 10:50 | 0:05 |
| 17 | light :: blue flicker | 3.5s : 0.5s on:off blue light | 10:50 | 11:30 | 0:40 |
| 18 | light :: red flash | solid red light (5 s) | 11:30 | 11:35 | 0:05 |
| 19 | light :: UV flicker | 3.5s : 0.5s on:off UV light | 11:35 | 12:15 | 0:40 |
| 20 | light :: red flash | solid red light (5 s) | 12:15 | 12:20 | 0:05 |
| 21 | light :: violet flicker | 3.5s : 0.5s on:off violet light | 12:20 | 13:00 | 0:40 |
| 22 | audio :: audio train | ring sound (1.2 s) repeated (45 s); 30 s gap; 1 final sound | 13:00 | 14:30 | 1:30 |
| 23 | light :: stripes | red, blue, red, UV, red, violet (4.5 s each with 500 ms gap) | 14:30 | 15:10 | 0:40 |
| 24 | audio :: thump | underwater thump sound (1.045 s; 1 copy) | 15:10 | 15:20 | 0:10 |
| 25 | light :: solid violet | solid violet light | 15:20 | 16:05 | 0:45 |
| 26 | audio :: soft taps | solenoid tap on felt every 3 s | 16:05 | 16:35 | 0:30 |
| 27 | audio :: hard taps | “soft" solenoid tap on stage every 3 s | 16:35 | 17:00 | 0:45 |

Numbers in parentheses signify the duration for which the stimulus is active. *1x*, *2x*, and *4x* denote 1, 2, and 4 copies of the stimulus (separated by small gaps).

**Table S6. Sound pressure levels measurements, C-weighted.**

| Condition | mean | S.D. |
|---|---|---|
| ambient | 63.4 | 1.2 |
| ‘ring’ | 90.6 | 0.8 |
| ‘drums’ | 97.4 | 0.4 |

Recordings are in a 1 s window around the peak. Recorded using a BAFX3608 digital sound level meter (BAFX Products). S.D., standard deviation.

**Table S7. Identifiers for referenced data.**

| Experiment | Run IDs |
|---|---|
| concentration–response | 6883,6885,6887,6888,6894,7029,7030,7031 |
| optimal-concentration | 7327,7329,7330,7331,7349,7473,7521,7522,7605 |
| NT-650 | 7667,7638,7706,7649,7971,7944,7697,7705,7672,7693,7975,7701,7666,7972,7710,7681,7974,7694,7686,7709,7679, 7682,7687,7731,7743,7983,7751,7987,7970,7771,7759,7980,7783,7762,7940,7957,7772,7801,7806,7824,7823,7977,7831, 7984,7828,8232,8233,8231,8238,8234,8226,8225,7943,8235,8237,8239,8241,7988,8240,8229,8230,8228,8227,8236,7833, 7832,7829,7830,7836,7834,7849,7842,7841,7847,7840,7850,7882,7894,7900,7895 |

IDs correspond to `runs.id` in the database.

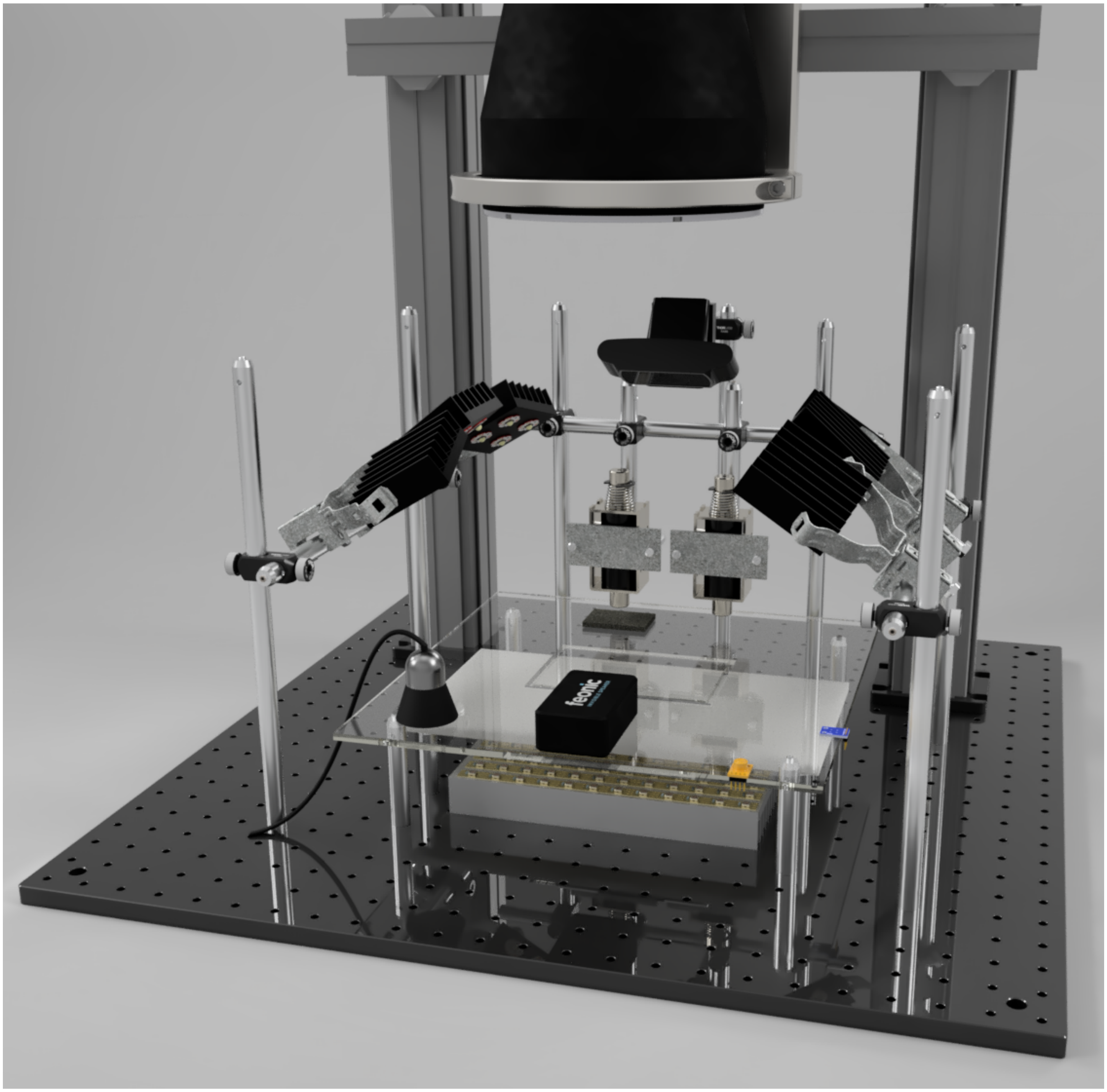


**Fig. S1.** 3D figure. Close-up of stage.

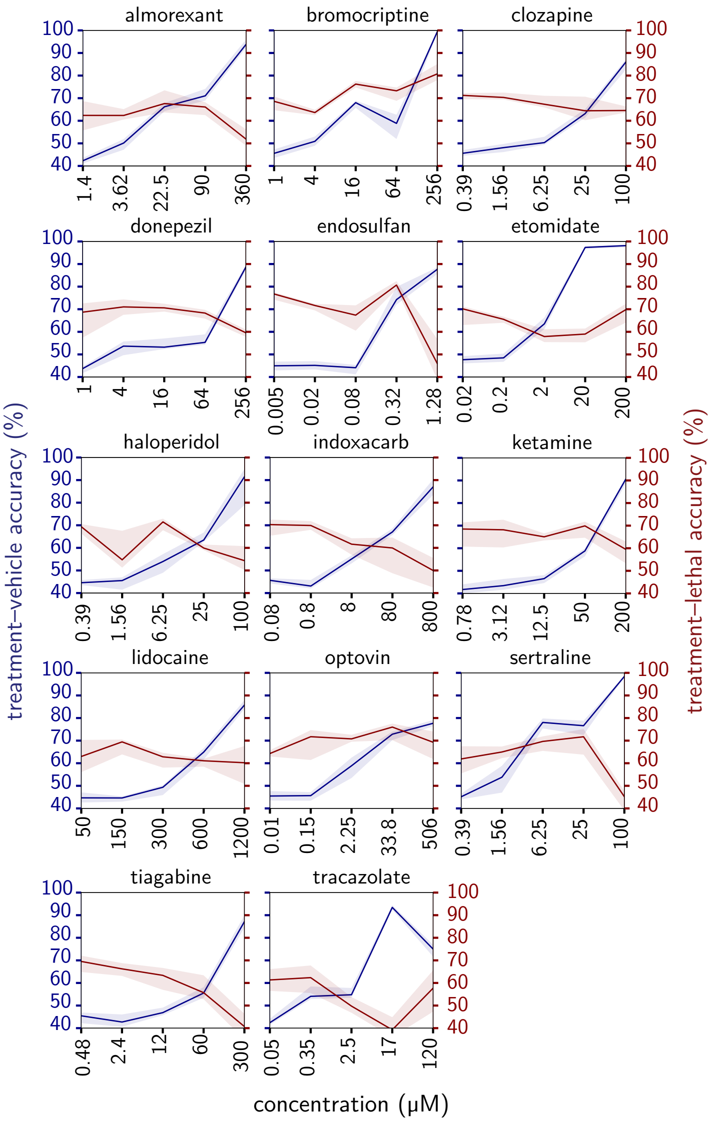


**Fig. S2.** Concentration–response for all 14 quality–control compounds.

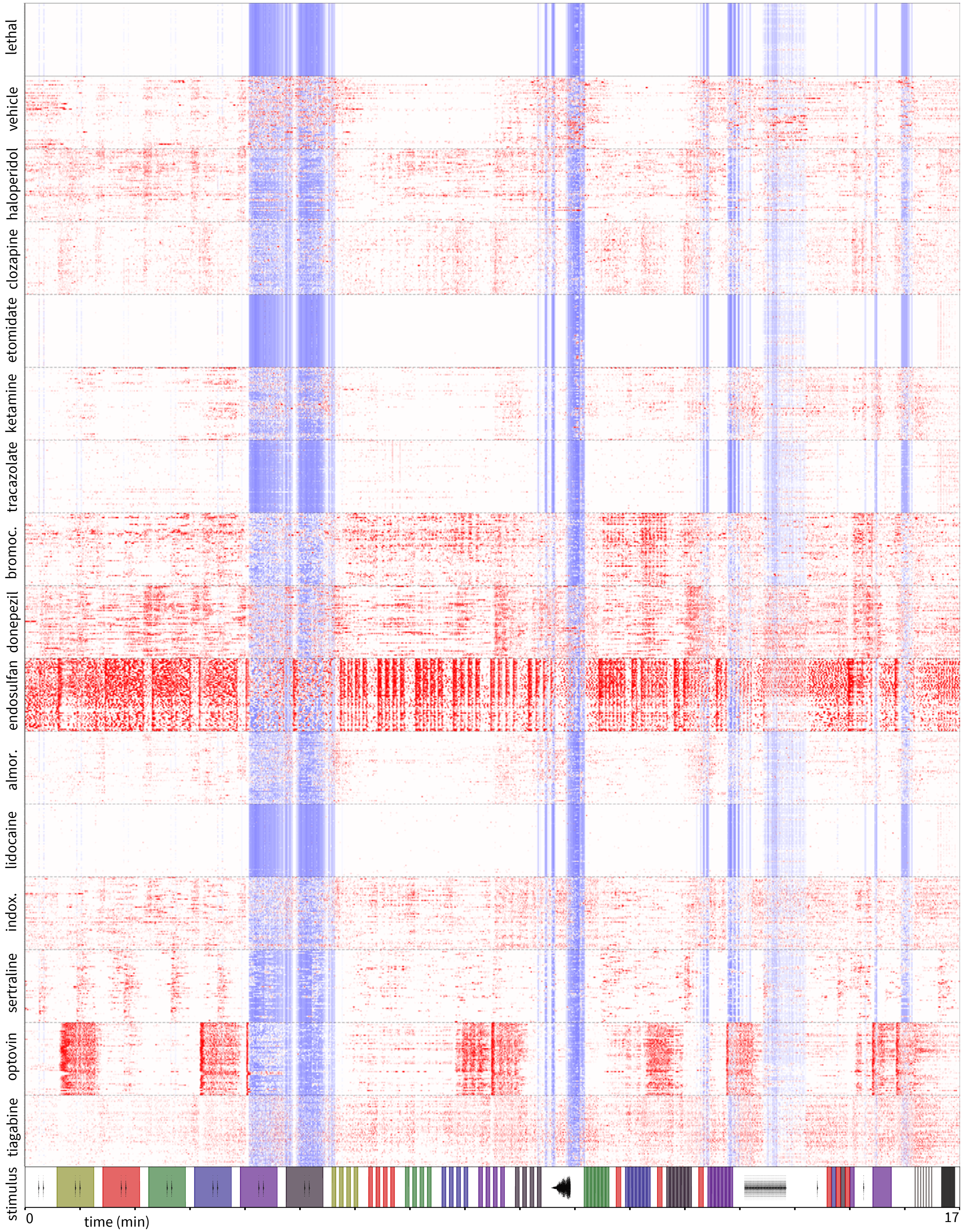


**Fig. S3.** Per-well motion traces of optimal-concentration data shown as a heatmap of Z-scores with respect to the mean of solvent controls. Values within 1 standard deviation were set to white. bromocr., bromocriptine; almor., alomorexant; indox., indoxicarb.

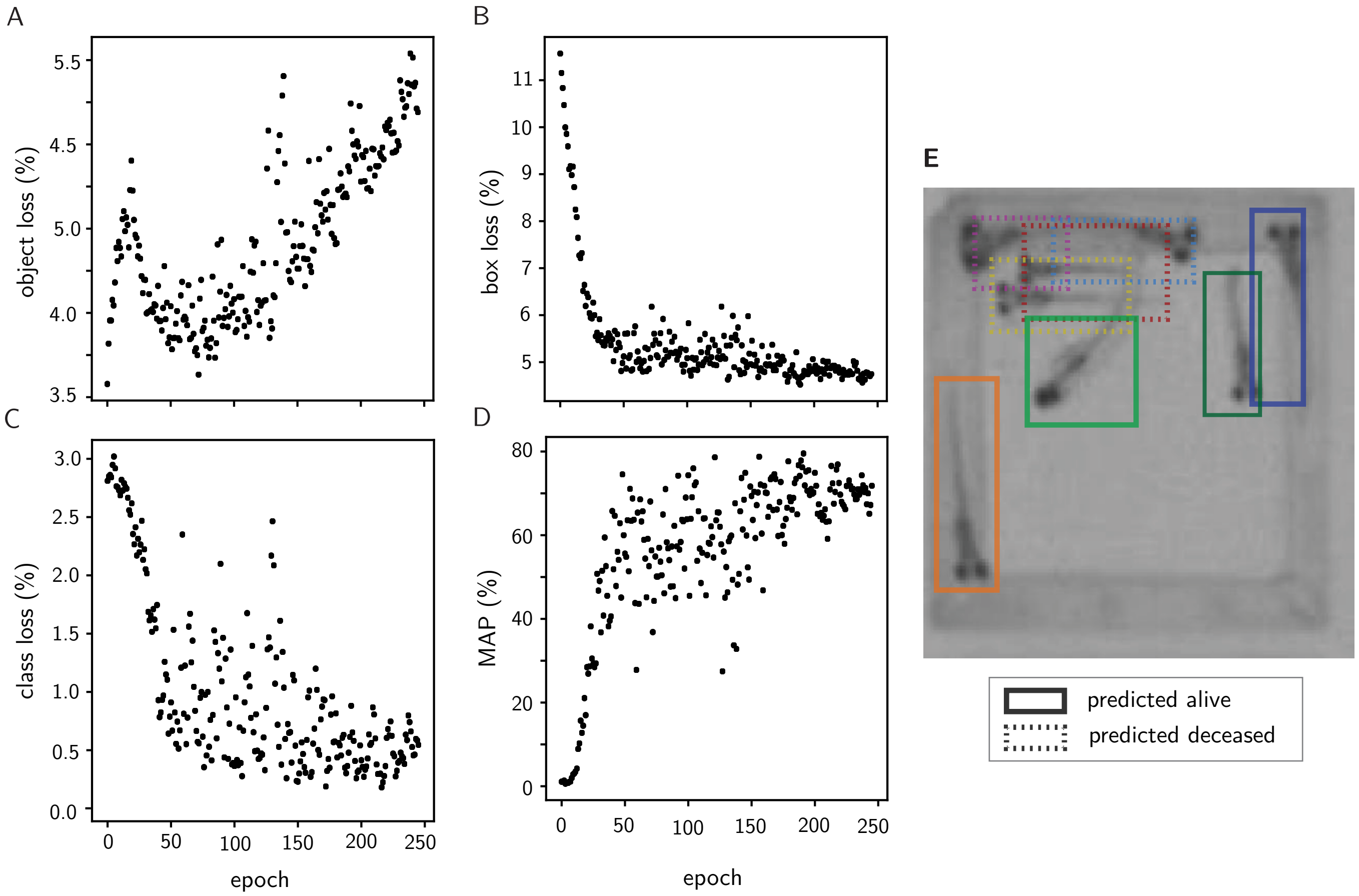


**Fig. S4.** Results for the You Only Look Once models. (**A–C:**) v alidation set loss curves for object (*A*), bounding box (*B*), classification of alive versus deceased (*C*). (**D**) Mean average precision (MAP) in validation set for intersection over union 0.5. (**E**) Example well with detected objects and labels.

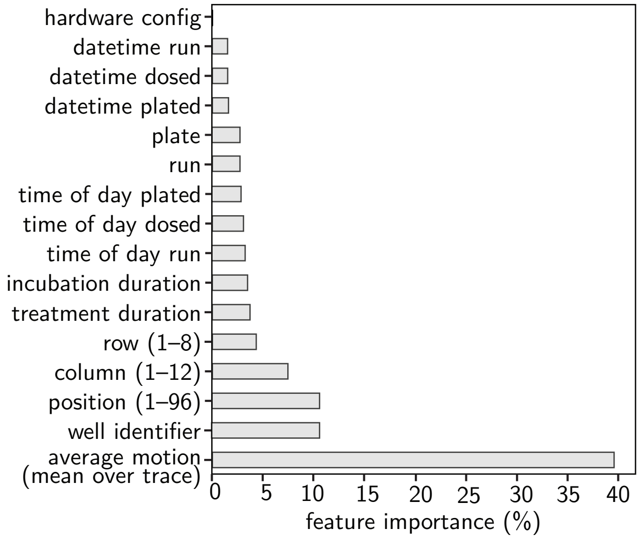


**Fig. S5.** Feature importance from a multiclass classifier trained on optimal-concentration quality–control compounds. For these experiments, *run* and *plate* are identical. *Incubation duration* refers to the duration between plating and dosing.


Douglas Myers-Turnbull, Jack C Taylor, Cole Helsell, Matthew N McCarroll, Chris S Ki, Tia A Tummino, Shreya Ravikumar, Reid Kinser, Leo Gendelev, Rebekah Alexander, Michael J Keiser, David Kokel

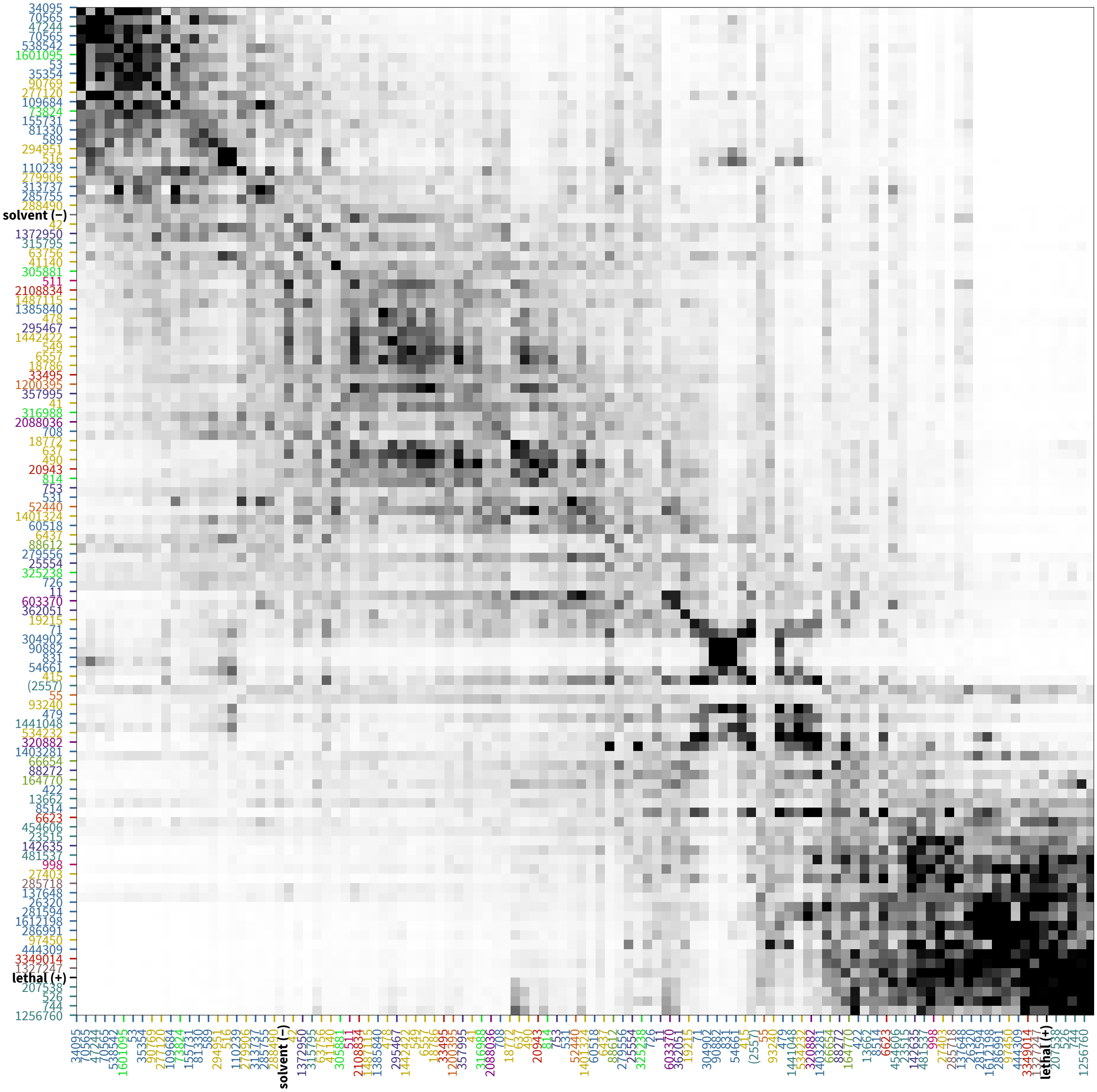


**Fig. S6.** Confusion matrix for NT-650 multiclass classification, sorted using confusion matrix ordering. Colors range from the 2nd percentile (white) to 98th percentile (black).

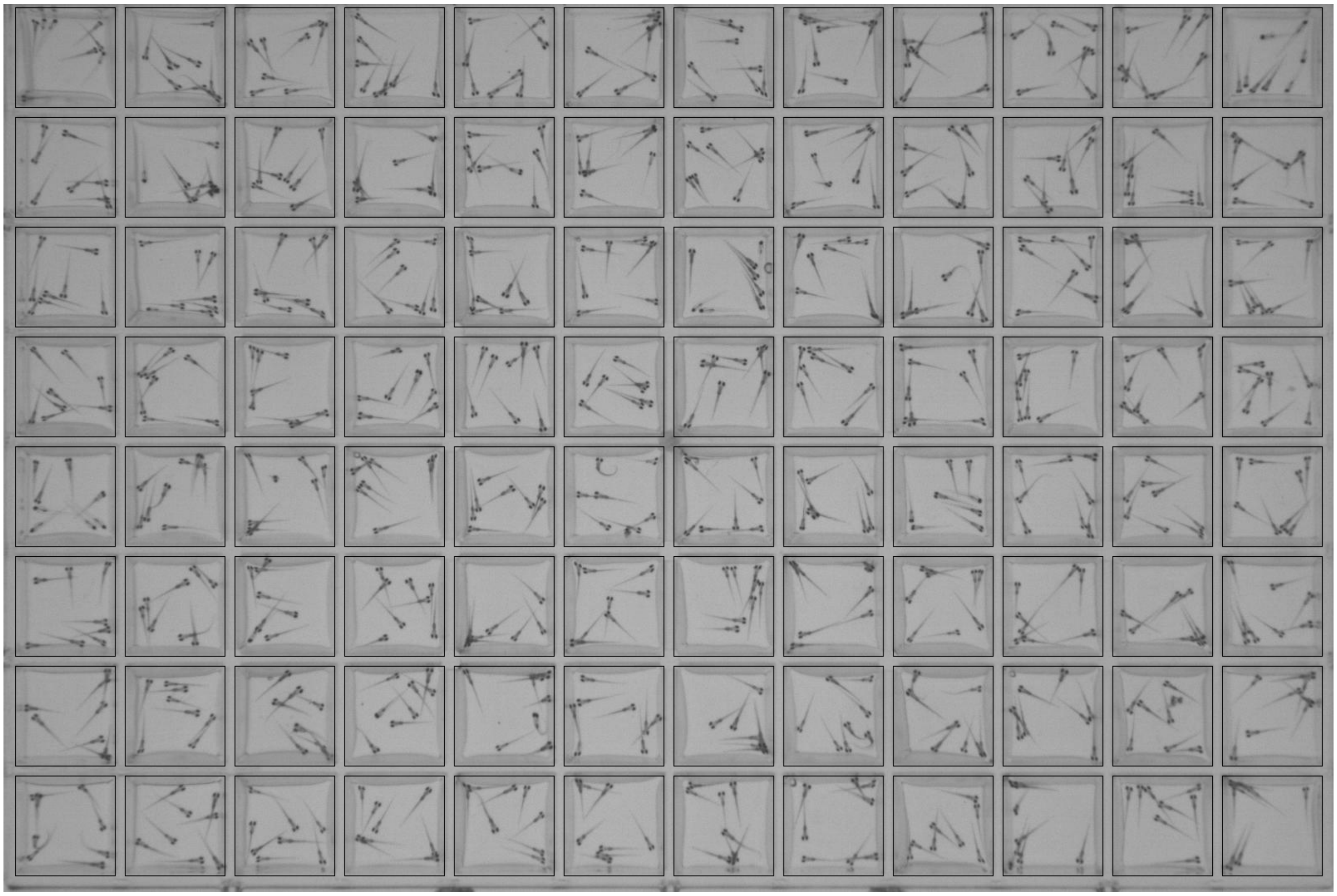

**Fig. S7.** Example captured frame with ROI overlay.

**Audio File S1. Ring stimulus.**

**Audio File S2. Thump stimulus.**

**Audio File S3. Drums stimulus.**

**Movie S1. Responses to 355 nm and 400 nm light. Quality is reduced to limit file size.**

**Movie S2. Exact-quality HEVC video in a 10 s no-stimulus assay.**
